# Modulation of visual cortex by hippocampal signals

**DOI:** 10.1101/586917

**Authors:** Julien Fournier, Aman B Saleem, E Mika Diamanti, Miles J Wells, Kenneth D Harris, Matteo Carandini

**Author notes:** equal contributions (Julien Fournier, Aman B Saleem, E Mika Diamanti).

## Abstract

Neurons in primary visual cortex (V1) are influenced by the animal’s position in the environment and encode positions that correlate with those encoded by hippocampus (CA1). Might V1’s encoding of spatial positions be inherited from hippocampal regions? If so, it should depend on non-visual factors that affect the encoding of position in hippocampus, such as the physical distance traveled and the phase of theta oscillations. We recorded V1 and CA1 neurons while mice ran through a virtual corridor and confirmed these predictions. Spatial representations in V1 and CA1 were correlated even in the absence of visual cues. Moreover, similar to CA1 place cells, the spatial responses of V1 neurons were influenced by the physical distance traveled and the phase of hippocampal theta oscillations. These results reveal a modulation of cortical sensory processing by non-sensory estimates of position that might originate in hippocampal regions.

---

There is increasing evidence that the responses of neurons in primary visual cortex (V1) are influenced by the animal’s position in the environment (Fiser et al., 2016; Haggerty and Ji, 2015; Ji and Wilson, 2007; Pakan et al., 2018; Saleem et al., 2018). Positions encoded by V1 neurons match those encoded by regions of the navigational system such as the hippocampus (Haggerty and Ji, 2015; Saleem et al., 2018) and fluctuations in the spatial position encoded by the V1 population correlate with those observed in hippocampus CA1 (Saleem et al., 2018). These correlated positional signals might reflect a propagation of sensory estimates from the visual system to the navigational system; alternatively, they might reflect an influence of positional estimates encoded by the navigational system, on visual processing.

If V1 responses are influenced by the navigational system, they should be modulated by the non-visual factors that affect the representation of space in hippocampus, and should correlate with hippocampal responses even in the absence of visual signals. Hippocampal place cells do not rely only on vision to encode the animal’s position; their place field is present also in the dark (Muller and Kubie, 1987; O’Keefe and Speakman, 1987); their spatial selectivity is influenced by the distance traveled in the environment (Chen et al., 2013, 2019; Gothard et al., 1996; Jayakumar et al., 2019; Ravassard et al., 2013) and on shorter timescales, their preferred place of firing depends on the phase of the ongoing 6-9 Hz theta oscillations (theta precession, O’Keefe and Recce, 1993; Skaggs et al., 1996). Here we asked whether similar non-visual factors affect the representation of space in V1.

## Results

To relate the spatial responses in area V1 to those of hippocampal area CA1, we recorded from both regions simultaneously while mice performed a spatial task in virtual reality (Figure **1A-1C**). Head-fixed mice ran on a wheel to explore a virtual corridor (Harvey et al., 2009; Saleem et al., 2018) defined by three landmarks (L1-L3; Figure **1A**). The corridor was semicircular and repeated into a full circle without interruption, with occasional periods of blank screen every 10-30 trials. We trained mice to lick in a position centered around landmark L1 for a water reward. To encourage the mice to use strategies beyond the discrimination of visual textures, the texture at landmark L1 alternated between the textures shown at L2 and L3 (a plaid and a grating, Figure **1A**). After ~6-8 weeks, mice licked exclusively when approaching the reward zone in more than 80% of the trials (Figure **1B**). We then used silicon probes to record from CA1 and V1 simultaneously (Figure **1C**). We focused on neurons that significantly modulated their firing rate across positions in the corridor (p < 0.01): 1,422 neurons in CA1 (out of 54 recording sessions, after excluding putative interneurons) and 1,109 neurons in V1 (out of 35 recording sessions). We identified putative interneurons in V1 based on the width of their spike waveform (21.7% of neurons; Figure **S1A**) (Bartho et al., 2004; Sirota et al., 2008). Other neurons are likely to be mostly pyramidal. Neurons in both V1 and CA1 responded similarly regardless of the texture shown in the reward position (grating or plaid), so we pooled all trials together (Figure **S1B-S1E**).

**Figure 1.**
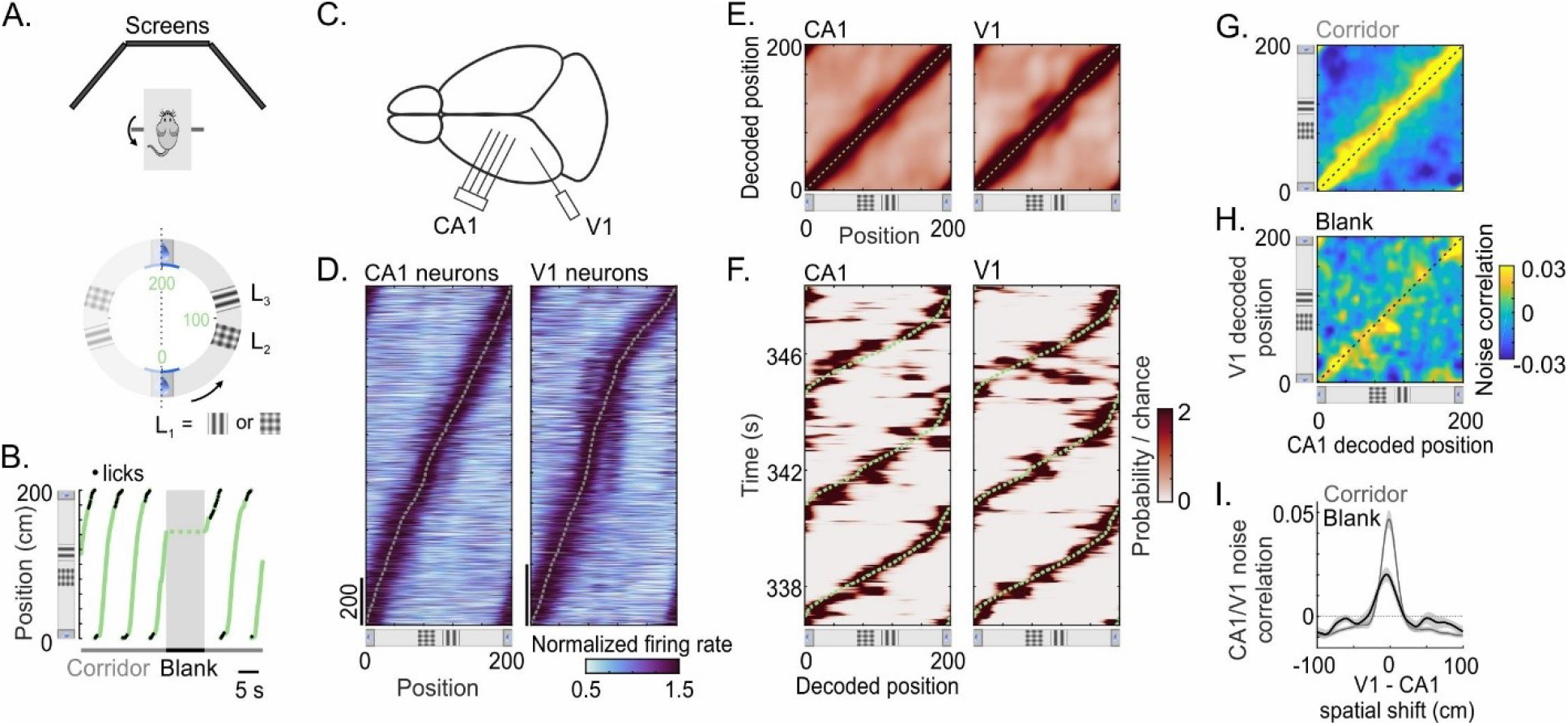
Spatial representations in V1 and CA1 correlate in the absence of visual cues. **A**. Schematic of the setup showing the running wheel surrounded by three computer screens (*top*) and the virtual corridor with three landmarks (L1, L2 and L3, *bottom*). **B**. Example trajectories, showing the locations of licks (*black dots*) and a blank interval when the task was temporarily interrupted by a gray screen. **C**. Schematic of the recordings with 32-electrode silicon probes in the CA1 region of the hippocampus and in primary visual area V1. **D**. Responses of neurons in CA1 (*left*) and V1 (*right*) that showed a significant modulation of firing rate across positions in the corridor (p < 0.01). Neurons are ordered by the position of maximal firing (*dotted curve*). For each neuron, the response profile was normalized by the mean firing rate across all positions. **E**. Average decoded probability of the animal’s position as a function of its actual position. The decoded probability was estimated for each recording session, from neurons recorded in CA1 (*left*, n = 42 sessions) or V1 (*right*, n = 33 sessions), and averaged across sessions. Color scale as in **F. F**. Decoded probability distribution estimated for three successive trials in one example session (CA1, *left*; V1, *right*). *Dotted curve*: actual position of the animal. **G**. Noise correlations between CA1 and V1 decoded probability measured for each decoded position when the mice were in the virtual corridor, averaged across recording sessions (n = 27 sessions with simultaneous recording). **H**. Same as **G**. during blank periods. **I**. Average noise correlations (± s.e.m) between CA1 and V1 decoded probability distributions, computed for different spatial shifts between the two distributions (CA1 – V1 spatial shift, n = 27 sessions). *Gray curve*: noise correlations measured in the corridor. *Black curve*: noise correlations measured during blank periods.

### Spatial representations in V1 and CA1 correlate in the absence of visual cues

Populations in both CA1 and V1 represented the spatial position of the animal in the corridor, and made correlated errors (Figure **1D-1G**; Saleem et al., 2018). Similar to CA1, V1 neurons had response profiles that tiled the entire corridor (Figure **1D**). Accordingly, we could use either area to decode the animal’s position with high accuracy (Figure **1E**). Nonetheless, there were occasional errors in the spatial representation of both V1 and CA1 (Figure **1F**), which could not be explained by position or speed. As previously shown (Saleem et al., 2018), these trial-by-trial fluctuations were spatially aligned with each other and significantly correlated (p < 0.05, n = 27 sessions; Figure **1G** and **1I**, *gray curve*).

Correlated fluctuations in the position encoded by V1 and CA1 could arise for two reasons. First, the computations estimating spatial position from visual input might be noisy, resulting in common fluctuations arising purely from visual processing. Second, the fluctuations might reflect an effect of non-visual factors on both areas, for example feedback of position signals from hippocampus to visual cortex, or a common effect of a third area on both.

Our data supported a role of non-visual factors: correlations in the position encoded by V1 and CA1 fluctuations were observed also in the absence of visual cues (Figure **1H-1I**). Every 10-30 trials, the virtual environment was interrupted by a blank screen (uniform gray) ~50 cm before the reward region. This blank period lasted 8.5 ± 2.5 s (s.d.) after which the animal resumed the trial in the same position as before the blank (Figure **1B**). Even during the blank periods, there was a significant correlation between the fluctuations in position decoded from V1 and CA1 (p < 0.05; Figure **1I**, *black curve*). This correlation was significant at p < 0.05 in 19/27 of recording sessions (p = 3.6 10^−25^, Fisher’s combined probability test), and it scaled with the correlation measured in the corridor (Pearson’s correlation coefficient = 0.48, p = 0.011; Figure **S2A**). During the blank periods, positions decoded from CA1 or V1 were unrelated to the distance run by the animal or the time passed; instead they fluctuated across a range of positions, with a bias towards the start of the corridor or the position where the animal was before the blank (Figure **S2C-S2F**). Correlations between V1 and CA1 decoded probabilities were present across a range of decoded positions (Figure **1H** and **S2B**). The correlation between V1 and CA1 could also be seen in pairs of CA1 and V1 neurons, both in the corridor and during the blank periods (Figure **S2G**).

### Spatial representations in V1 and CA1 are influenced by distance run

Hippocampal spatial representations are affected by a number of non-visual cues, amongst which idiothetic information (i.e. the distance traveled in the environment) is fundamental (Campbell et al., 2018; Chen et al., 2013, 2019; Gothard et al., 1996; Jayakumar et al., 2019; Ravassard et al., 2013). We next asked whether idiothetic cues could also affect representations of space in visual cortex.

To investigate the effect of idiothetic cues, we changed the gain of the wheel by ±20% in a fraction of trials (43% ± 15 s.d.), so that the animal had to run a longer or shorter distance than usual to reach a given visual position (Figure **2A-2C**). This manipulation affected behavior: mice tended to lick earlier when the distance was longer (low gain), and later when the distance was shorter (high gain, Figure **2B**). Consistent with a dominant role of vision in this task, the shift in licking position was smaller than would be expected if the mice were solely counting steps (Figure **2B**). Nevertheless, the shift was substantial (low gain, −3.1 cm ± 2.4 s.d.; high gain, 2.4 cm ± 2.2 s.d.) and consistent across sessions (Figure **2C**).

**Figure 2.**
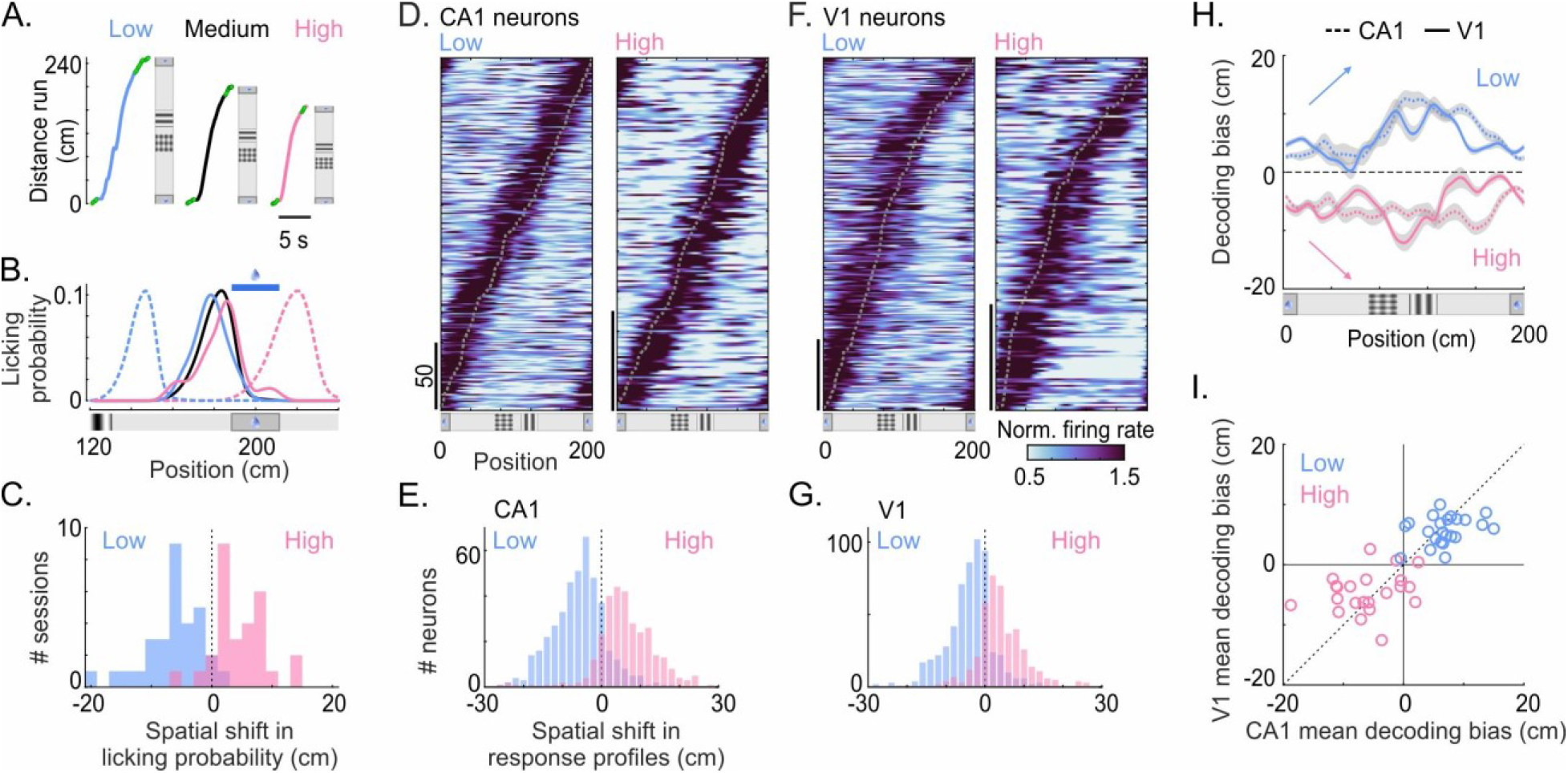
Spatial representations in V1 and CA1 are influenced by distance run. **A**. Examples of single-trial trajectories for low, medium and high gain trials, showing the effect of gain on physical distance run. **B**. Licking probability as a function of the animal’s position on medium (*black*), low (*blue*) and high (*pink*) gain trials for all correct trials in one example session. *Dashed curves* show the prediction if licks were based on physical distance run from the previous reward zone (40-cm shift). **C**. Spatial shift in licking probability measured from low or high gain trials across sessions, relative to the mean licking position on medium gain trials. **D**. Response profiles of CA1 neurons that showed a significant spatial shift of their response (in any direction) on low (*left*) or high (*right*) gain trials (low gain; 53.9% of 421 neurons; high gain; 54.8% of 303 neurons; p < 0.05). Response profiles were ordered according to the position of maximal firing on medium gain trials (*dotted curve*). For each neuron, the response profile was normalized by the mean firing across all positions. **E**. Distribution of spatial shifts in CA1 response profiles at low (*blue*) or high (*pink*) gain, using the response profile at medium gain as a reference. All neurons which had a significant modulation of their firing rate across positions of the corridor were included in this distribution. **F-G**. Same as in **D-E** for V1 neurons (low gain; 39.6% of 566 neurons; high gain; 37.9% of 403 neurons, p < 0.05). **H**. Decoding bias (± s.e.m) measured on low (*blue*) or high (*pink*) gain trials from V1 (*solid line*) or CA1 (*dashed line*). *Arrows* indicate the bias that would accumulate if spatial representation depended only on physical distance run. **I**. Comparison of the mean decoding bias (averaged across positions) between V1 and CA1 (n = 27 sessions).

Having established that the physical distance run influenced the animal’s decisions to lick, we asked whether it also influenced the activity of CA1 and V1 neurons (Figure **2D-2G**). To compare responses across gain conditions, we focused on neurons whose firing rates were significantly modulated across positions in the corridor on both medium and low or high gain trials (CA1: low gain, 421/1422 neurons; high gain, 303/1422; V1: low gain, 566/1109 neurons; high gain, 403/1109; p < 0.01). Changes in gain did not affect the mean firing rate of V1 or CA1 neurons (Figure **S3A** and **S3B**), but did affect the position of their neural responses relative to the position of the visual cues. As expected (Chen et al., 2013; Gothard et al., 1996; Jayakumar et al., 2019), CA1 place cells tended to fire at earlier positions at low gain, when the distance run was longer (53.9% of 421 neurons, p < 0.05) and at later positions at high gain, when the distance run was shorter (54.8% of 303 neurons, p < 0.05) (Figure **2D,2E** and **S3C**). Many V1 neurons exhibited a similar behavior: even though the landmarks were encountered at the same visual positions in all conditions, the neurons fired earlier on the track when the distance was longer (low gain; 39.6% of 566 neurons, p < 0.05) and later when the distance was shorter (high gain; 37.9% of 403 neurons, p < 0.05, Figure **2F,2G** and **S3D**). This spatial shift affected V1 neurons responding throughout the corridor, regardless of their preferred position (Figure **2F** and **S3D**). Responses of putative V1 interneurons and putative pyramidal cells shifted by similar amounts (low gain: p = 0.20; high gain: p = 0.07, rank sum test).

Accordingly, changing the gain shifted the spatial representations encoded by populations in both CA1 and V1 (Figure **2H-2I**). CA1 and V1 tended to encode a position ahead of the animal at low gain (when the distance run was longer) and behind the animal at high gain (when the distance run was shorter; Figure **2H**, **S3E** and **S3F**). These effects could not be explained by changes in running speed, visual speed or eye position (Figure **S3G-S3L**) or by an interaction of the latency of visual responses in V1 with the speed of the virtual environment (Figure **S3M** and **S3N**). Across recording sessions, areas V1 and CA1 showed a consistent decoding bias at low and high gain (Figure **2I**). We conclude that idiothetic signals modulate spatial coding in V1 similarly to how they modulate spatial coding in CA1.

### V1 neurons are modulated by CA1 theta oscillations

We next asked whether V1 neurons are influenced by another non-visual factor that affects place cells: the 6-9 Hz theta oscillation (Figure **3**). As expected (Buzsáki et al., 1983), the theta oscillation modulated the firing rate of most CA1 place cells (86.6%, n = 1,422 neurons, p < 0.05; Figure **3A** and **3B**), with firing rates highest near the trough of the theta cycle (i.e. 180°; p = 10^−181^, Rayleigh test, Figure **3C**). Hippocampal theta oscillations can also entrain neurons in somatosensory and prefrontal cortices (Jones and Wilson, 2005; Sirota et al., 2008; Zielinski et al., 2019). We observed a similar effect in V1, where the hippocampal theta oscillation significantly modulated the firing of 23.4% of V1 neurons (p < 0.05, n = 1,109; p = 6.0 10^−153^, Fisher’s combined probability test; Figure **3D** and **3E**). The theta modulation of firing rates was weaker in V1 than in CA1 (p = 10^−76^, rank sum test). V1 neurons modulated by the theta oscillation had diverse preferences for positions in the corridor (Figure **S4A**). Similar to previous observations in somatosensory and prefrontal cortices (Sirota et al., 2008), in V1 the theta oscillation modulated narrow-spiking putative interneurons more than putative pyramidal neurons (36.9% vs. 19.6%; Figure **3E**). Putative pyramidal neurons varied in their preference for theta phase, whereas putative interneurons tended to prefer the peak of the theta cycle (i.e. 0°; p = 0.036, Rayleigh test; Figure **3F**). These data demonstrate that V1 activity is significantly coupled to the same theta oscillation as CA1 place cells.

**Figure 3.**
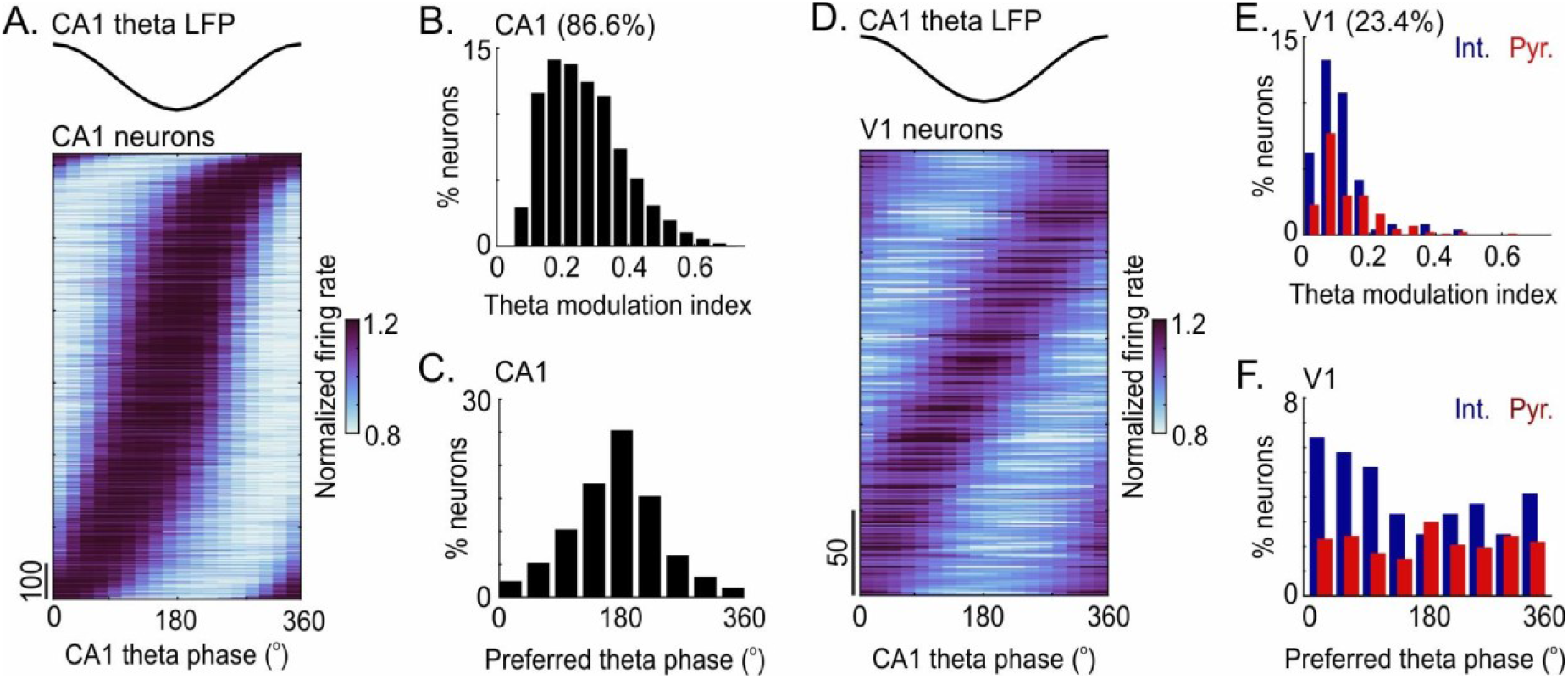
V1 neurons are modulated by CA1 theta oscillations. **A**. Firing rate of CA1 neurons as a function of the phase of theta oscillations recorded in hippocampus (CA1 theta LFP). Each neuron’s firing rate was normalized by the mean firing across phases. Neurons were ordered according to phase of maximal firing. Only neurons with a significant modulation of their firing rate across theta phases are represented (86.6%, n = 1,422 neurons; p < 0.05). **B**. Distribution of the amplitude of theta modulation (*Theta modulation index*) for CA1 neurons shown in A. **C**. Distribution of the theta phase of maximal firing (*Preferred theta phase*) for CA1 neurons shown in A. **D-F**. Same as **A-C**, showing the activity of V1 neurons relative to the theta oscillation measured in CA1. 23.5% of V1 neurons showed a significant modulation of firing rate across theta phase (n = 1,109 neurons; p < 0.05, **D**). Histograms in **E** and **F** distinguish putative interneurons (*Int., blue*; n = 240) and pyramidal cells (*Pyr., red*; n = 869), identified from their spike waveform. Percentages are relative to the total number within each class.

### Spatial representations in V1 and CA1 oscillate with theta phase

Our CA1 recordings showed typical theta phase precession in hippocampal place cells (O’Keefe and Recce, 1993; Ravassard et al., 2013; Figure **4A-4E**). Indeed, CA1 neurons fired ahead of their preferred position at late phases of the theta cycle (180° to 360°), and behind their preferred position at early phases (0° to 180°, Figure **4A-4C**). This spatial oscillation in the position of the place field with theta phase was seen in 48.5% of CA1 neurons (p < 0.05). The spatial oscillation typically went from negative to positive at phases near 180° (phase precession, Figure **4D**) so the population encoded a position ahead of the animal at late phases and behind the animal at early phases (Figure **4E**).

**Figure 4.**
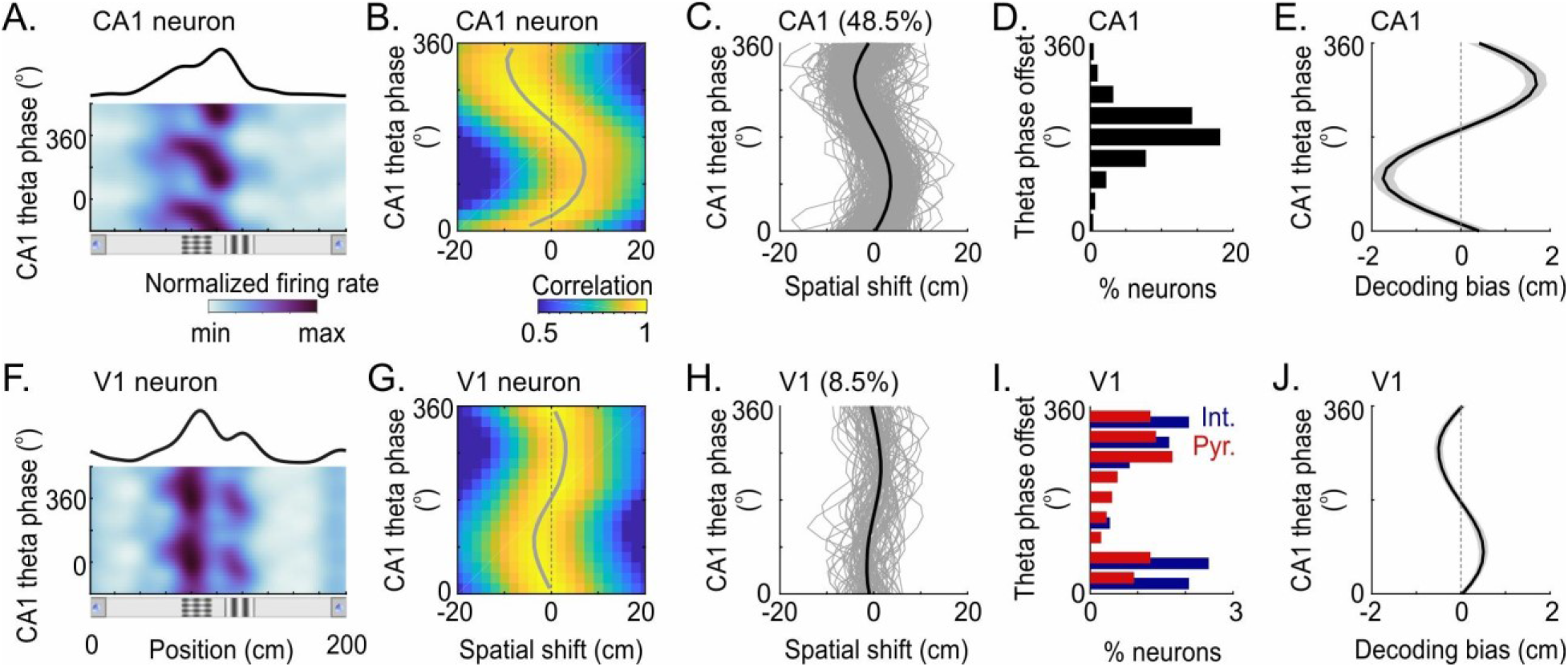
Spatial representations in V1 and CA1 depend on the phase of theta oscillation. **A**. Firing rate of a CA1 neuron example as a function of position (*x* axis) and phase of CA1 theta oscillations (*y* axis). *Black curve*: spatial response profile of the same neuron. **B**. Spatial cross-correlogram between the response profiles at each theta phase and the average response profile, for the place cell example shown in **A**. *Gray curve*: position of maximal correlation across theta phases. **C**. Spatial shift in response profiles across theta phases (*gray curve* in **B**) for CA1 neurons with a significant spatial shift across theta phases (*gray curves*; 48.5%, n = 1,422 neurons; p < 0.05; *Black*: average). **D**. Distribution of the theta phase offset (point of zero-crossing when the spatial shift went from negative to positive), for CA1 neurons with a significant spatial shift across theta phases. Percentages are expressed relative to the total number of CA1 neurons (n = 1,422 neurons). **E**. Mean bias in the Bayesian probability decoded from entire populations of CA1 neurons, as a function of the phase of the CA1 theta oscillation. *Gray band* shows ± s.e.m. (n = 42 sessions). **F-G**. Same as in **A-B** for a V1 neuron example recorded simultaneously to the CA1 neuron shown in **A. H-I**. Same as **C-D** for V1 neurons with a significant spatial shift across theta phases (8.5%, n = 1,109 neurons; p < 0.05). Histograms in **I** distinguish putative interneurons (*Int., blue*; n = 240) and pyramidal cells (*Pyr., red*; n = 869). Percentages are relative to the total number within each class. **J**. Same as in **E**, calculated from entire populations of neurons in V1 (n = 33 sessions).

Intriguingly, positions encoded by V1 neurons also depended on the phase of the theta oscillation measured in CA1 (Figure **4F-4J**). A significant oscillation in the position of the response profile across theta phases was seen in 8.5% of V1 neurons (p < 0.05, n = 1109; Fisher’s combined probability test, p = 2.4 10^−17^; putative pyramidal cells, 8.2% of 868; putative interneurons, 9.5% of 241; Figure **4F-4H** and **S4C**). The phase of this spatial oscillation varied widely across V1 neurons, but tended to go from negative to positive at phases near 0/360° (i.e. opposite to the typical phase precession of CA1 place cells; putative pyramidal cells, p = 1.4 10^−4^; putative interneurons, p = 7.7 10^−4^; Rayleigh test; Figure **4I**). V1 neurons that significantly shifted their response across the theta cycle were found throughout the corridor (Figure **S4B**). This dependence of V1 spatial selectivity on theta phase was visible not only in a fraction of V1 neurons but also in the population as a whole: positions decoded from entire populations of V1 neurons were shifted ahead of the animal at late theta phases and behind at early theta phases (Figure **4J**). Hippocampal theta oscillations thus appear to modulate V1 neurons not only in terms of firing rate but also in terms of spatial selectivity. Albeit smaller than in CA1 (Figure **4C,4H** and **S4D-S4E**), this modulation echoes the phase precession observed in hippocampus.

## Discussion

We found that the information encoded in primary visual cortex correlates with that in the hippocampus even without visual input, and that it is influenced by non-visual factors that are known to contribute to spatial coding in hippocampus: the physical distance run and the hippocampal theta oscillation. These results suggest that signals from the navigational system may shape visual processing in cortical areas.

Our observation that the activity of V1 neurons is coupled to hippocampal theta oscillations extends a number of previous observations. Theta phase locking of visual neurons has been observed in monkey V4 during a working-memory task (Lee et al., 2005) and in local field potential recordings from rats (Zold and Hussain Shuler, 2015). Hippocampal theta oscillations, moreover, have been shown to entrain the activity of cortical neurons in prefrontal, and somatosensory areas (Jones and Wilson, 2005; Sirota et al., 2008; Zielinski et al., 2019).

We found that positions encoded by V1 neurons shifted spatially across phases of a theta cycle. Theta phase precession has been observed beyond hippocampus, e.g. in entorhinal cortex (Hafting et al., 2008; Jeewajee et al., 2014), ventral striatum (van der Meer and Redish, 2011) and prefrontal cortex (Jones and Wilson, 2005; Zielinski et al., 2019). Our data show that theta phase coding may also affect primary sensory areas that receive no direct input from hippocampus. It is unclear whether the theta phase modulation observed in V1 is inherited or locally generated. Nonetheless, the fact that the phase of the spatial oscillation varied across V1 neurons and was more often opposite to the theta phase precession observed in dorsal hippocampus suggest that V1 may be influenced by different parts of the hippocampal network. Indeed, in hippocampus, the phase of the theta oscillation varies along the septotemporal axis (Lubenov and Siapas, 2009). V1 might thus be more influenced by ventral hippocampus or CA3 where theta phase precession is also 180° out of phase relative to dorsal CA1 (Lubenov and Siapas, 2009; Royer et al., 2010).

Our results show that in a familiar environment, V1 is influenced by the physical distance run: when the distance is increased or decreased, V1 neurons respond at a position that is intermediate between the actual position where visual cues are displayed and the physical distance at which the animal used to encounter these visual cues. A possibly similar effect was seen in V1 neurons selective to a rewarded position; their firing may depend on the physical distance run, at least when no salient visual cue is present (Pakan et al., 2018). Here, we showed that this integration of visual cues and distance information in V1 may also happen throughout the trajectory of the animal in the corridor: in our dataset, V1 neurons exhibited a shift in their response when the distance changed, whether they fired preferentially at the reward position or elsewhere.

The network of connections that could explain a top-down influence of the navigation system on visual cortex is unknown. Several intermediate cortical areas may be involved, such as the anterior cingulate cortex (Fiser et al., 2016), or retrosplenial cortex (Mao et al., 2017; Vélez-Fort et al., 2018). Feedback signals from the navigation system might also influence visual processing in thalamic neurons upstream to V1 (Busse, 2018; Hok et al., 2018). A promising way to probe these connections may be during sleep, when V1 and CA1 tend to activate together, potentially replaying sequences of activation observed during active exploration (Ji and Wilson, 2007).

Why should the spatial representation in V1 be modulated by these non-visual signals? We suggest that V1 is part of a large network that combines multiple streams of input to transform sensory information from self-centered to world-centered coordinates and estimate the animal’s current spatial location (Nau et al., 2018). Although activity in V1 will have a visual bias compared to other regions, it will combine an estimate of visual position based on sensory evidence and an estimate of position based on non-visual cues such as the distance traveled from a previous visual location. The presence of self-motion information in V1 may thus reflect top-down signals carrying a prediction of the visual scene (Fiser et al., 2016; Poort et al., 2015), which would synergize with dynamics imposed by external sensory inputs to generate a more accurate positional estimate.

## Acknowledgements

We thank C. Reddy for help with surgeries and C. Barry for discussions. This work was funded by Human Frontier Science Program (fellowship LT001022/2012-L to J.F.) and EC Horizon 2020 (fellowship 709030 to J.F.), by Wellcome Trust/Royal Society (Sir Henry Dale Fellowship 200501 to A.B.S.), by EPSRC (PhD award F500351/1351 to E.M.D.), by the Wellcome Trust (grant 205093 to M.C. and K.D.H.), and by the Simons Collaboration on the Global Brain (grant 325512 to M.C. and K.D.H.). M.C. holds the GlaxoSmithKline/Fight for Sight Chair in Visual Neuroscience.

## Author contributions

Conceptualization: J.F., A.B.S., K.D.H., M.C. Investigation: J.F., A.B.S. and M.J.W. Analysis: J.F., A.B.S., E.M.D. Writing: J.F., A.B.S., E.M.D., K.D.H., M.C. Funding acquisition: J.F., A.B.S., E.M.D., K.D.H., M.C. Supervision: M.C.

## Methods

All experimental procedures were performed in accordance with the UK Animal Scientific Procedure Act 1986, under project and personal licenses issued by the UK Home Office.

### Surgical procedure

Data were collected from ten C57BL/6J male mice. The surgical procedure is similar to that described previously (Saleem et al., 2018). In brief, mice were implanted on their left hemisphere with a 4-mm diameter chamber at 4-10 weeks of age, under deep isoflurane anesthesia. Mice were left to recover for 3 days during which they received anti-inflammatory drug (Carprofen/Rymadil, oral administration). After recovery, mice were water-restricted and moved to light-shifted conditions (9 a.m. light off, 9 p.m. light on). Mice were then trained once a day during their dark cycle, approximately at the same hour of the day, for several weeks. After they reached sufficient performance in the task, we performed two 1-mm craniotomies under deep isoflurane anesthesia: one over CA1 (1.0 mm lateral, 2.0 mm anterior from lambda), and the other over V1 (2.5 mm lateral, 0.5 mm anterior from lambda). Recordings were carried on the subsequent 5-8 days with one recording session per day. Between recordings, the chamber was covered with silicon (KwikCast, World Precision Instrument).

### Electrophysiological recordings

Recordings were performed with multi-site silicon probes connected to a Blackrock amplifier sampling at 30 kHz. Neurons in the dorsal CA1 region of the hippocampus were recorded using a silicon probe with 32 electrodes arranged in 8 tetrodes spread over four shanks (2 tetrodes per shank spaced by 150 μm vertically, 200-μm distance between shanks, Neuronexus A4×2-tet). The pyramidal layer of CA1 was detected by the increase in power of theta oscillations (6-9 Hz) and the large number of detected units. Neurons in primary visual cortex were recorded using a 32-electrode linear probe (20 μm electrode pitch; Neuronexus A1×32-Edge). In V1, the probe was inserted so that the most superficial electrode was ~150 μm under the cortical surface. V1 and CA1 were recorded simultaneously in 35 sessions.

The broadband signal was high-pass filtered at 500 Hz, and spike-sorted using Klustakwik and Klustaviewa (Rossant et al., 2016). We identified 2748 neurons in CA1 (out of 54 recording sessions) and 1,433 neurons in V1 (out of 35 recording sessions). Hippocampal interneurons were identified based on the duration of their spike waveform (through to next peak < 600 *μ*s) and the shape of their spike time auto-correlogram (no prominent peak between 3-8 ms) (Chen et al., 2013). All identified interneurons were excluded from further analysis. Single units in V1 were classified as putative interneurons if the duration of their spike waveform was < 600 *μ*s from trough to next peak (Figure **S1A**).

### Local field potential and theta oscillations

Hippocampal local field potential (LFP) was extracted by filtering the broad-band signal between 0.1 Hz and 100 Hz. The LFP signal from the tetrode with the largest number of pyramidal neurons was filtered between 6 Hz and 9 Hz to obtain the theta-band oscillation. The instantaneous phase of the theta oscillation was measured by detecting the peaks in the oscillation and measuring the relative time from one peak to the next, which we then converted into phase. Peaks which occurred earlier than 60 ms after the preceding one were discarded. Theta phase values were centered so that the overall firing rate of the population of CA1 pyramidal neurons peaked at 180° theta phase.

### Virtual environment and behavior

Mice navigated a virtual environment by walking on a cylindrical wheel made of polystyrene (Saleem et al., 2018). The movement of the wheel was measured with a rotary encoder (2400 pulses per rotation, Kübler, Germany). Distance traveled on the wheel was used to translate the virtual environment presented on three LCD monitors (Hanns-G, 60Hz refresh rate) placed in front of the animal at 34-cm viewing distance. To correct for luminance drop-off, monitors were covered with Fresnel lenses. The eye position and pupil size were monitored with an infrared camera (DMK21BU04.H, Imaging Source) and a zoom lens (MVL7000, Navitar) at 25 Hz. Eye tracking was performed offline by fitting an ellipse to the pupil and measuring the center of mass and the area of this ellipse (Stringer et al., 2019).

The virtual environment was a circular track made of two identical 200-cm semicircular corridors (Figure **1A**). The main advantage of using a continuous circular track is that it avoids edge-effects at the start and the end of the animal’s trajectory. The track was covered with a 16.7-cm periodic grating. A wall ran along the right side of the track and was adorned with a periodic Gaussian-filtered white noise which repeated every 16.7 cm. The left side of the track had no wall. Three landmarks (L1, L2, L3) made of 24-cm long cylindrical tunnels (16 cm in diameter) were placed along the corridor and centered at 0 cm (L1), 83 cm (L2) and 117 cm (L3). The inside surface of these tunnels was covered with either a vertical grating (L1, L3) or a plaid (L1, L2). The outside surface of the tunnels was covered with a horizontal grating. Mice could navigate this environment indefinitely and there was no visual discontinuity going from one semicircular corridor to the next one. Navigation was however interrupted every 10 to 30 trials with a blank screen of several seconds (8.5 s ± 2.5 s.d.). This blank screen always occurred ~50 cm before the reward zone. Mice usually continued running during the blank period but withheld licking until the task resumed. After the blank period, navigation resumed in the same position as before the blank, regardless of the distance that the animal run during the blank screen.

Mice were trained to lick selectively in a region centered around landmark L1 (± 12 cm). Licks were detected using a custom-made infrared detector. When licks were detected in the correct region, the animal was rewarded with ~2μL water by opening of a pinch valve (NResearch, USA). To ensure that the animal did not learn the task by simply discriminating the visual texture associated to landmark L1, the visual texture displayed at landmark L1 alternated between a grating and a plaid every other trial.

Mice usually tended to lick in bouts that we detected by identifying succession of licks that were spaced by less than 20 cm. We then labelled licks as being correct or incorrect depending on whether they were part of a bout of licks that overlapped with the reward region. The licking probability distribution was computed from the spatial position of the first lick of these bouts, considering only correct licks. We measured lick counts and occupancy as a function of position. The lick count map and the occupancy map were circularly smoothed with a Gaussian window (8-cm s.d.). Licking profiles were defined as the ratio between the lick count map and the occupancy map. To calculate licking probability across position, the resulting distribution was normalized to sum to 1. Mice typically learned this task in ~6-8 weeks, i.e. they eventually licked exclusively when approaching the reward region and nowhere else in more than 80% of the trials. Once the animal had learned the task, we changed the gain of the virtual environment on a fraction of trials (43% ± 15 s.d.). On trials where the gain was lower (0.8), mice had to run 250 cm to reach the reward zone; on trials where the gain was higher (1.2), they had to run 166 cm to the reward zone. Gain changes were made in blocks of 5-10 trials. This manipulation was typically introduced 2-3 days before recording. Mice performed correctly on more than 70% of the trials when the gain was low or high.

We only considered time points when the animal run speed was >5 cm.s^−1^, except for estimating the licking probability; trials where incorrect licks were detected were also excluded from our analyses.

### Spatial response profiles

Spike times were resampled at the screen refresh rate (60 Hz) and the position of the animal was binned into 2-cm bins. Trials were sorted according to the gain of the virtual environment and response profiles were computed separately on medium, low and high gain trials. The spatial response profile of each neuron was computed by measuring spike counts and occupancy as a function of position. The spike count map and the occupancy map were circularly smoothed with a Gaussian window (8-cm s.d.). Spatial response profiles were defined as the ratio of the smoothed spike count map over the smoothed occupancy map.

To test for the significance of response profiles, we circularly shifted in time each neuron spike train 500 times by a random period > 5 s. Neurons were considered to have a significant modulation of their firing rate by position if the maximal amplitude of their response profile was higher than the 99-percentile of the distribution of response amplitude computed from the shuffled controls. Neurons were then ordered according to their preferred position, defined as the position of maximal firing rate.

To estimate the shift in response profiles across gain conditions, we computed the spatial cross-correlogram between medium and low- or high-gain response profiles, for neurons which had a significant response modulation across compared gain conditions (p < 0.01). The spatial shift was defined as the position of the peak in the spatial cross-correlogram between response profiles. To test for the significance of the spatial shift, we shuffled 500 times the gain values associated to the spikes of each neuron. Neurons were considered to have a significant shift of their response profile on low or high gain trials when this shift was larger than the 95-percentile of the distribution of shifts measured from the shuffled controls.

We also measured mean firing rates as a function of gain. To test if changes in gain significantly affected mean firing rates, we circularly shifted in time each neuron’s spike train 500 times. The mean firing rate was considered to be significantly different between medium and low or high gain trials, if the measured difference in firing rate was higher than the 99-percentile of the distribution of the same difference computed from the shuffled controls.

We also computed the response profile of each neuron as a function of position along the two successive semicircular corridors, which only differed by the presence of a grating or a plaid at landmark L1 (Figure **1A** and **S1B-S1C**). We measured the difference in peak firing rate between the second and the first corridor at matching positions (i.e. 200 cm apart). To test if the difference in peak firing rate was significant, we shuffled 500 times the identity of the semicircular corridor (first vs. second). A neuron was considered to respond differently in the first and second corridor if the difference in peak firing rate was larger than the 99-percentile of the distribution of the same difference computed from the shuffled controls.

### Theta modulation

Firing rate as a function of the phase of theta oscillations was computed similar to spatial response profiles. Theta phases were binned into 20° bins. We measured the spike counts and occupancy as a function of the phase of theta oscillations. The spike count map and the occupancy map were circularly smoothed with a Gaussian window (40° s.d.). The average firing rate across theta phases was defined as the ratio of the smoothed spike count map over the smoothed occupancy map. The preferred theta phase of firing was measured for each neuron from the position of the peak. The theta modulation index was computed as

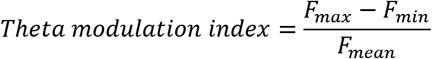

Where *F_max_* and *F_min_* correspond to the maximum and minimum firing rate across theta phases; *F_mean_* is the mean firing rate across all theta phases.

To measure the significance of theta modulation of firing rate, we circularly shifted in time each neuron’s spike train 500 times by a random period > 5 s. Firing rate was considered significantly modulated by theta phase if its theta modulation index was higher than the 95-percentile of the distribution of theta modulation index computed from the shuffled controls.

### Theta phase dependent response profiles

To assess how spatial selectivity changed as a function of theta phase, we measured average firing rate as a function of both the position of the animal and the phase of theta oscillations. Positions were binned in 2-cm bins and theta phase into 20° bins. Spike count and occupancy were measured for each position and theta phase bin. The spike count map and the occupancy map were circularly smoothed using a 2d-Gaussian window (40° × 8-cm s.d.). The *theta phase* × *position* response profile was defined as the ratio between the spike count and occupancy maps. We then measured the spatial cross-correlogram between the response profile estimated at each theta phase with the mean response profile averaged across all phases. The spatial drift of the response profile across a theta cycle was defined as the maximum of this cross-correlogram across theta phases. This maximum position curve was fitted with a sinusoid from which we measured the theta phase offset (i.e. the phase of zero-crossing) and the amplitude. To measure the significance of the spatial drift across theta phases, we shuffled the theta phases of the spikes of each neuron 500 times. Neurons were considered to have a significant spatial drift when the amplitude of the drift measured from the spatial cross-correlogram was higher than the 95-percentile of the distribution of spatial drifts measured from the shuffled controls. The spatial cross-correlogram has two main advantages over the more classical linear-circular correlation (Kempter et al., 2012): 1) it is independent of the static modulation of firing rate by theta phases and 2) it does not require the identification of the peak and extent of the place field. For comparison, we also estimated the relationship between the theta phase and position of spikes by estimating the linear-circular correlation in the *theta phase* × *position* maps (Kempter et al., 2012). With this metric, response profiles were considered to show a significant association between theta phase and position when their linear-circular correlation coefficient was smaller or larger than the 5- or 95-percentile of the distribution of correlation coefficients measured from the shuffled controls. Using this method, we found similar fractions of response profiles that were spatially shifted across phases of a theta cycle (Figure **S4E**).

### Bayesian decoding

To decode positions from populations of simultaneously recorded neurons, we used an independent Bayes decoder, assuming spike times followed a Poisson distribution. Decoding was performed for recording sessions where >10 neurons showed a significant modulation of their firing by position (CA1, n = 42 sessions; V1, n = 33 sessions; n = 27 sessions with simultaneous recording). For every time bin, we estimated the probability of the animal being at a given location (P(x|R), decoded probability) from the spike count of CA1 or V1 neurons, using the following formula (Zhang et al., 1998):

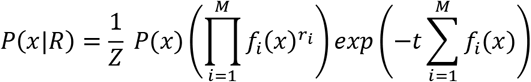

Where *x* is the position of the animal, *r_i_* is the spike count of cell *i, f_i_*(*x*) is the spatial response profile of cell *i* estimated as described above, *M* is the total number of neurons, *t* is the size of the spike counting window and *Z* is a normalizing constant that ensures that the resulting distribution sums to 1.

The response profiles *f_i_*(*x*) were computed from medium gain trials as described above. The spike count window was fixed to 250 ms (except when we assessed the bias in decoded probability across a theta cycle where we used 50 ms; Figure **4E** and **4J**). To avoid over-fitting, the posterior probability decoded from medium gain trials was computed using a 20-fold cross-validation procedure: response profiles were calculated from 95% of the data and the posterior probability was estimated from the spike counts recorded in the remaining 5%; this procedure was repeated iteratively on different subsets until all trials were decoded.

To minimize the impact of running speed on decoded probability, the running speed distribution measured from medium gain trials was divided into 5 quantiles at each position and the decoder was run for each of these speed ranges independently. For low and high gain conditions, data were sliced according to the same absolute speed ranges as in the medium gain condition. We used the average response profiles computed from medium gain trials to decode positions from the spike counts measured on low or high gain trials.

The mean decoded probability (Figure **1E** and **S3E-S3F**) was computed by averaging the posterior probability for each position of the mouse. The resulting distribution was smoothed with a 2d-Gaussian window (2 × 2-cm s.d.) and the mean decoded trajectory was estimated by computing the weighted circular average of this probability distribution for every position of the animal. To estimate the decoding bias between gain conditions, we measured the difference between the mean decoded trajectory computed in the low or high gain condition with the mean decoded trajectory computed in the medium gain condition. The standard error of this decoding bias was estimated using a 20-fold Jackknife resampling procedure.

### Noise correlations

Noise correlations in the virtual corridor were measured from correct trials that were not interrupted by a blank screen. To compute the noise cross-correlograms between V1 and CA1 decoded probability, we first computed the mean decoded probability for every position and speed bins (defined as described in **Bayesian decoding**). At every time point, we subtracted the mean decoded probability estimated at the corresponding position and speed of the animal from the probability distribution decoded at that time point. The noise cross-correlogram was finally obtained by computing the Pearson’s correlation between the residual CA1 and V1 distributions, after shifting circularly the CA1 distribution in space by various amounts. To test for the significance of the measured CA1-V1 noise correlation, V1 and CA1 decoding probability were shifted relative to one another 500 times by a random time period (> 5 s).

We considered that CA1 and V1 shared significant noise correlation in a particular recording session when the measured Pearson’s correlation coefficient was higher than the 95-percentile of correlation coefficients obtained from the shuffled controls.

To measure the noise correlation between pairs of individual CA1 and V1 neurons recorded simultaneously, we first subtracted from each neuron’s spiking activity the mean spike count corresponding to the position and speed of the animal (defined as in **Bayesian decoding**). The remaining spiking activity corresponded to the residual that could not be explained by position or speed. The noise correlation (or spike count correlation) between two neurons was estimated by computing the Pearson’s correlation between the residual spiking activity of these two neurons. To test for the significance of the measured spike count correlations, we shifted the neuron’s residual spike counts relative to one another by a random time period (> 5 s) 500 times. We considered two neurons to be significantly correlated when the measured spike count correlation was smaller or larger than the 5- or 95-percentile of spike count correlation coefficients obtained from the shuffled controls.

Noise correlations were also estimated over periods where the task was interrupted by a blank screen using a similar method. We first excluded the first and last 500 ms of the blank screen periods. In the rare case when mice licked during the blank, we also excluded a 500-ms time window centered around the lick. We then defined 5 speed ranges based on quantiles of the distribution of run speeds during blank screen periods. The noise correlation between CA1 and V1 decoded probability was computed within each speed range. At every time point, we subtracted the mean decoded probability estimated at the corresponding speed of the animal from the probability distribution decoded at that time point. We next obtained the noise cross-correlogram by computing the Pearson’s correlation between the residual CA1 and V1 distributions, after circularly shifting the CA1 distribution across space by various amounts. Similarly, to estimate the spike count correlations between pairs of neurons during blank periods, we subtracted the mean spike count corresponding to the speed of the animal at each time point. The spike count correlation between pairs of CA1 and V1 neurons was then measured as the Pearson’s correlation between their residual spiking activity. The significance of the correlation coefficients measured during the blank periods was estimated using the same method as described above for correlation coefficients measured in the corridor.

## Supplemental Information

**Figure S1.**
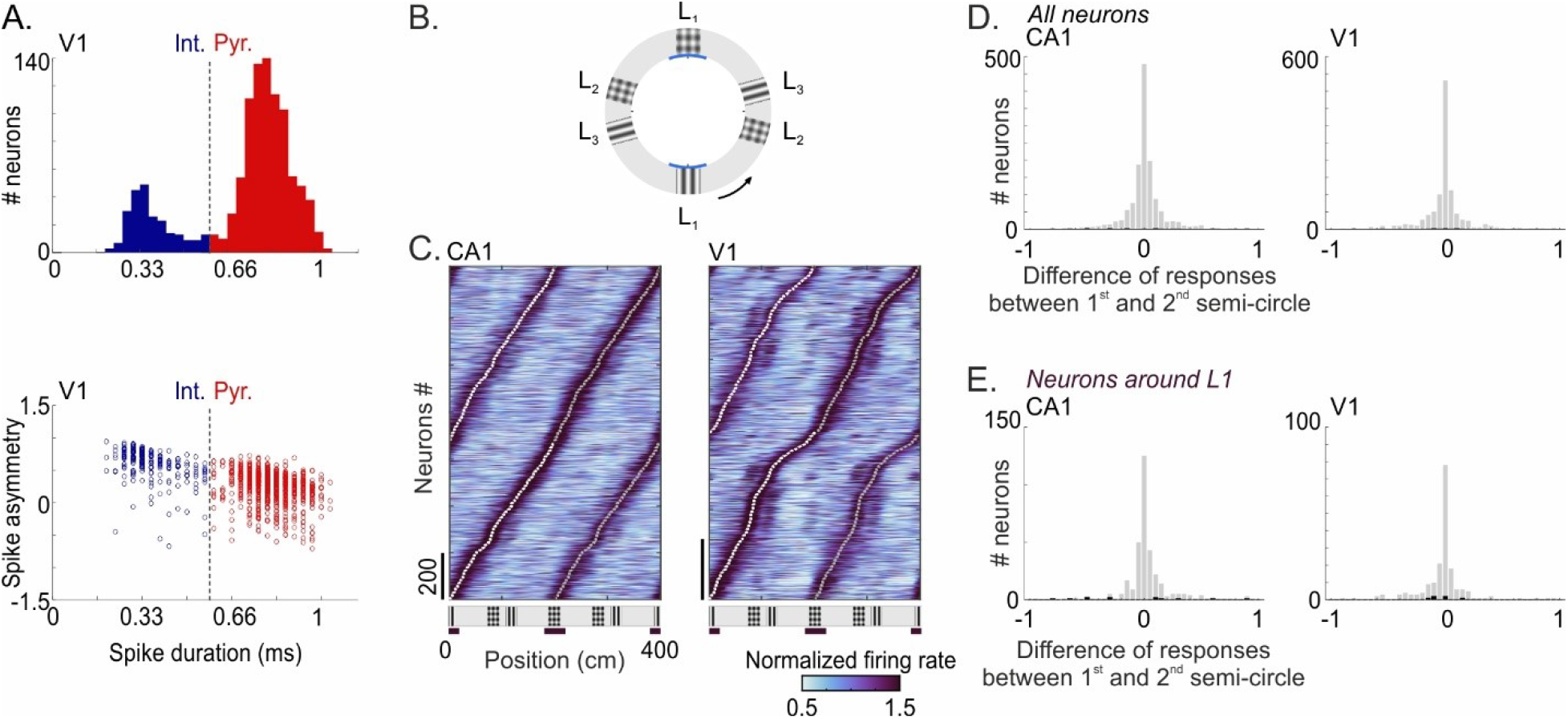
CA1 and V1 neural responses are similar whether the texture displayed at the reward location is a grating or a plaid. **A**. Putative neocortical interneurons and pyramidal cells were isolated from the duration of their extracellular spike waveform (Bartho et al., 2004; Sirota et al., 2008). *Top*, distribution of the duration of spike waveforms (trough to next peak; high-pass cutoff, 500 Hz) across V1 neurons (*Blue*, putative interneurons; *red*: putative pyramidal neurons; threshold: 0.60 ms). *Bottom*, other metrics such as the spike asymmetry (Sirota et al., 2008) did not help to better separate putative interneurons and pyramidal cells in our dataset. **B**. The virtual corridor was defined by three landmarks (L1, L2 and L3) and repeated in a full circle with texture at landmark L1 alternating between a grating and a plaid every other trial. **C**. Response profiles of CA1 (*left*) and V1 (*right*) neurons estimated as a function of position along the two successive semicircular corridors, which differed by the texture associated with landmark L1 (grating vs. plaid). Neurons are ordered according to the position of their peak response. Matching positions in the two semicircular corridors (200 cm away) are indicated by the dashed lines. Although the texture at the reward zone (L1) alternated between a grating and a plaid, responses were very similar between the two corridors. **D**. Distribution of the difference between peak responses in the first (starting with a grating, *white line* in B) and second (starting with a plaid, *gray line* in B) corridor, for CA1 (*left*) and V1 (*right*) neurons. The difference is expressed relative to the mean firing rate across all positions. *Black bars*: neurons with significant difference in peak firing rate between the first and second corridor (CA1, 1.19%, n = 1422 neurons; V1, 1.26%, n = 1109 neurons; p < 0.01). **E**. Same as **C** after selecting only neurons which had a maximal firing rate within 40 cm around landmarks L1 (neurons with significant difference in firing rate: CA1, 3.41%, n = 348 neurons; V1, 3.73%, n = 205 neurons; p < 0.01).

**Figure S2.**
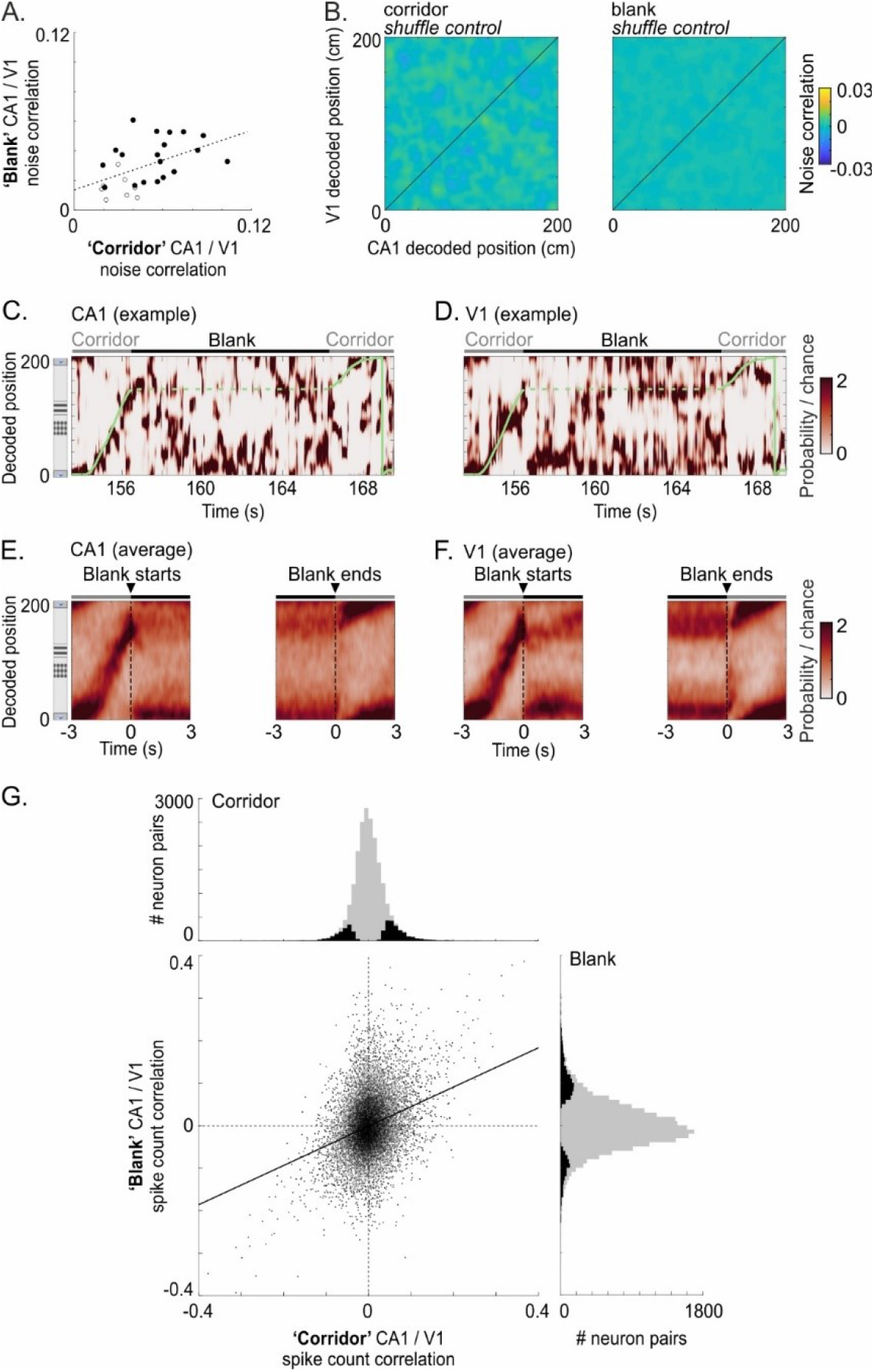
Correlations between V1 and CA1 representations in the absence of visual cues. **A**. Noise correlation between CA1 and V1 decoded probability distribution, measured either in the corridor (*x* axis) or during the blank periods (*y* axis). Each dot corresponds to one recording session (n = 27). In the corridor, all sessions showed a significant noise correlation between CA1 and V1 decoded probability (p < 0.05). In the blank periods, this correlation was significant in 70.4% of all sessions (p < 0.05; n = 19 out of 27, *filled circles*). The correlation measured in the corridor and during the blank periods scaled with each other (Pearson’s correlation coefficient = 0.48, p = 0.011). *Dotted line*: regression line. **B**. Noise correlations between V1 (*y* axis) and CA1 (*x* axis) decoded probability, measured for each decoded position independently, after shuffling time points within conditions. **C**. Bayesian probability decoded from CA1 for one trial example which was interrupted by a blank period. *Green line*: position of the animal. **D**. Bayesian probability decoded from V1 for the same trial as in **C. E**. Bayesian probability decoded from CA1 as a function of time from the start (*left*) or the end (*right*) of the blank period, averaged across all recording sessions (CA1, n = 42 sessions). **F**. Same as **E** for V1 (V1, n = 33 sessions). **G**. Spike count correlations between pairs of CA1 and V1 neurons recorded simultaneously (n = 20,563 pairs). Histograms show the distribution of spike count noise correlations in the corridor (*top*) or during the blanks (*right*), across all neuron pairs (*gray bars*). Black bars include only neuron pairs which had significant spike count correlation (p < 0.05; corridor, 17,8%; blank, 11.8%). The correlation coefficients measured during the blanks were significantly correlated with those measured in the corridor (Pearson’s correlation coefficient = 0.30; p < 0.002, permutation test; *Solid line* shows regression line).

**Figure S3.**
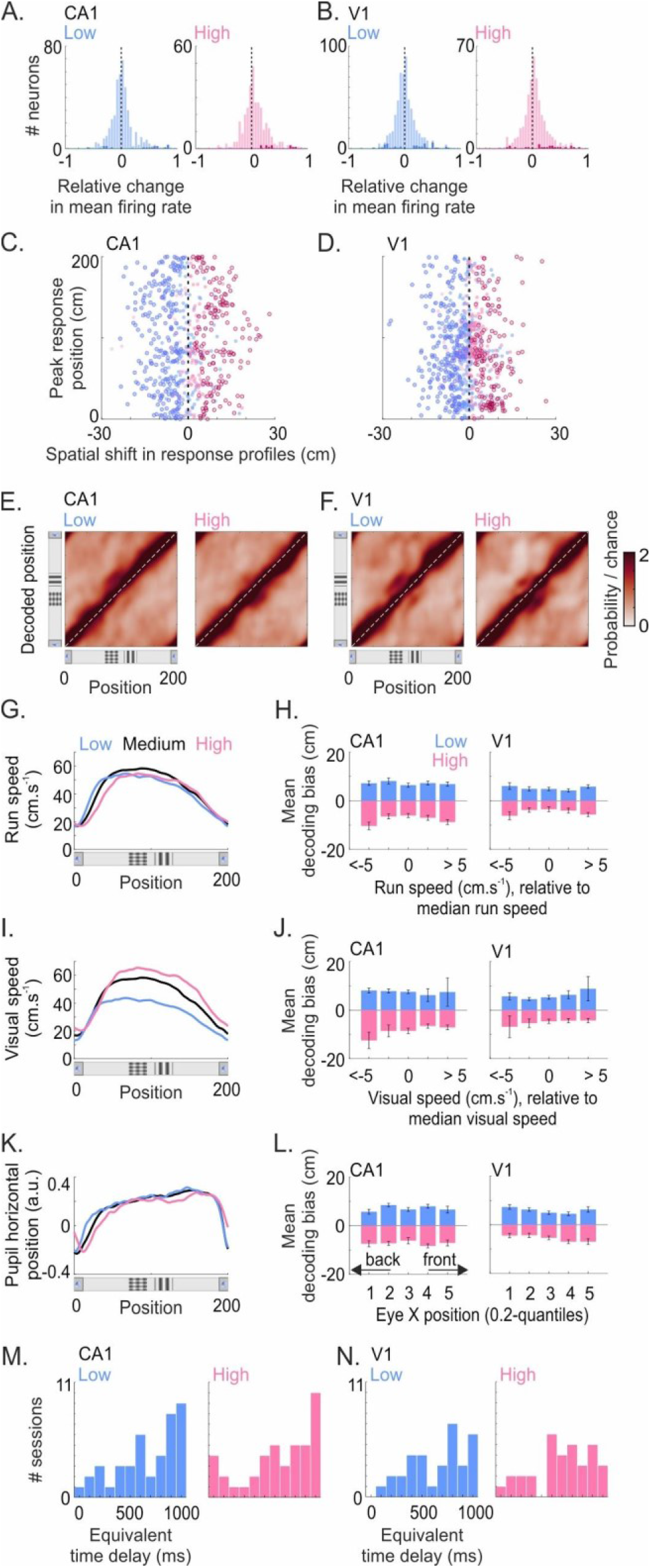
Positions encoded by V1 and CA1 are influenced by the physical distance run. **A**. Distribution of the difference in mean firing rate between medium gain trials and low (*left*) or high (*right*) gain trials, for CA1 neurons (n = 1422 neurons). The difference was normalized by the mean firing rate on medium gain trials. Darker colors represent neurons which had a significant change in firing rate on low or high gain trials compared to medium gain (low gain, 5.2%; high gain, 4.9%; p < 0.01). **B**. Same as A for V1 neurons (proportion of V1 neurons with significant change: low gain, 8.3%; high gain, 6.5%; p < 0.01). **C**. Spatial shift in CA1 response profiles on low (*blue*) and high (*pink*) gain trials, as a function of the position of the maximal firing along the corridor. Each dot corresponds to one neuron. Closed circles correspond to neurons with a significant spatial shift. **D**. Same as **C** for V1 neurons. **E**. Average Bayesian probability distribution decoded from CA1 on low (*left*) or high (*right*) gain trials. **F**. Same as **E** for V1. **G**. Mean running speed as a function of positions in the corridor. Running speed profiles were similar across gain conditions, except at the start of the corridor where mice used to accelerate slightly earlier or later. **H**. Mean decoding bias measured in CA1 (*left*) and V1 (*right*) for different ranges of running speed. Speed ranges were defined as the deviation of the running speed from the median of the running speeds at the animal’s current position on medium gain trials. **I-J**. Same as **G-H** for visual speed. Given similar running speed profiles across gain conditions (**I**), the visual speed was different between gain conditions. Speed ranges in **J** were defined as the deviation of the visual speed from the median of the visual speeds at the animal’s current position on medium gain trials. **K-L**. Same as **G-H** for pupil position along the horizontal axis. Pupil position was correlated with animal speed: gazing more forward when speeding up. The ranges of pupil position were defined by 0.2-quantiles of the distribution of eye positions at the animal’s current position on medium gain trials. **M**. Distribution of the time delays in neural responses that might explain the spatial decoding bias observed across CA1 recording sessions on low (median: 800 ms; *left*) or high (median: 750 ms; *right*) gain trials. Since mice ran at similar speeds across gain conditions, the virtual environment moved faster or slower depending on the gain (**I**). The spatial shifts in decoded positions could thus potentially reflect a temporal delay in the response of these neurons, which would translate into a displacement of their response in space at low or high gain. To estimate this ‘*equivalent time delay*’, we shifted the spike trains backward by various time delays, before performing again the same decoding analysis. The ‘equivalent time delay’ was measured as the time delay that minimized the decoding bias on low or high gain trials. **N**. Same as **M** for V1 recording sessions. The time delays that minimized the decoding bias measured in V1 (median: low gain, 750 ms; high gain, 650 ms) are not compatible with the latency of visual responses commonly observed in V1.

**Figure S4.**
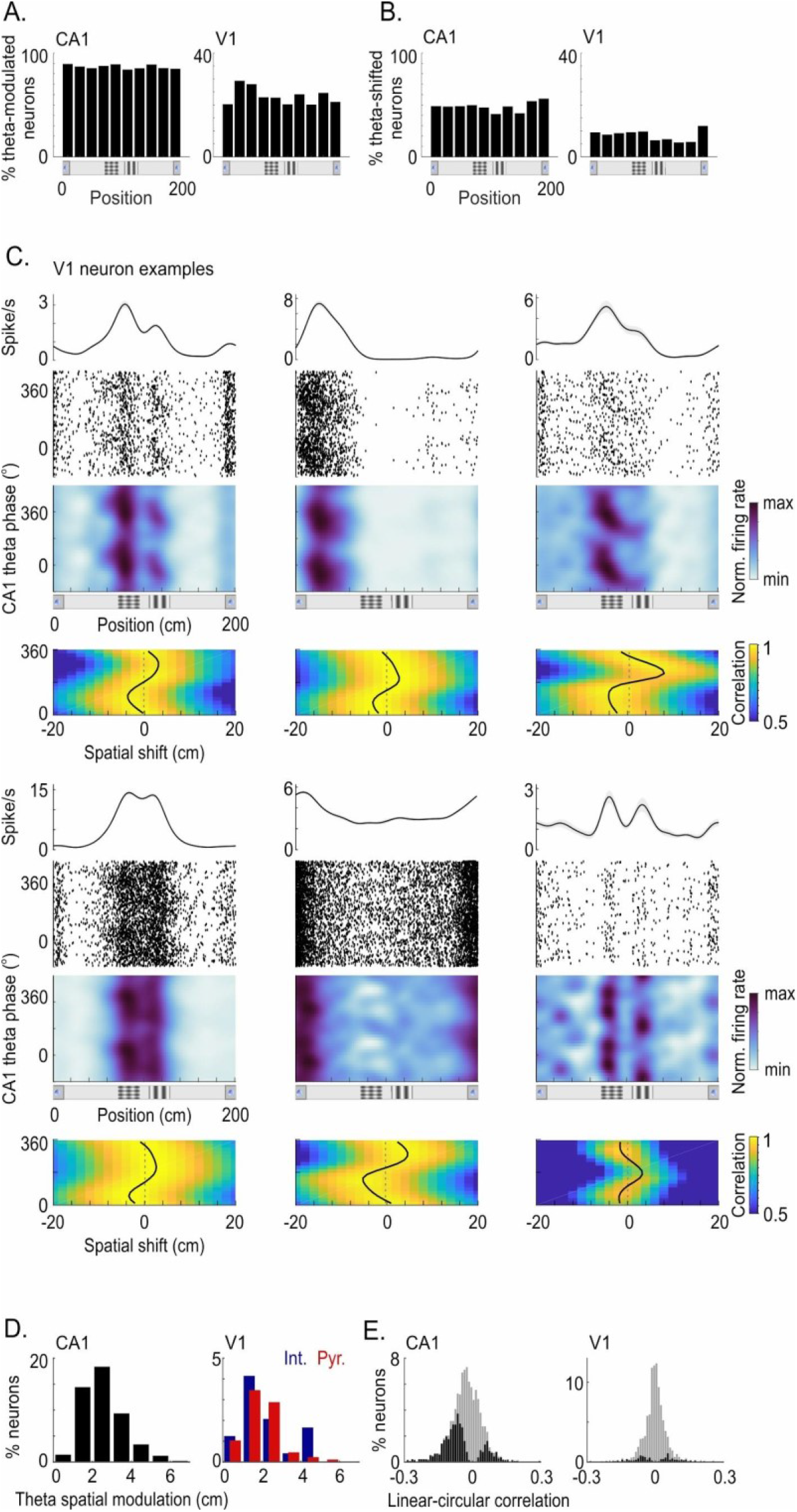
Theta phase dependence of V1 and CA1 responses. **A**. Percentages of CA1 (*left*) or V1 (*right*) neurons with a significant modulation of their firing rate by theta phases (‘*Theta-modulated neurons*’; p < 0.05), as a function of the position of maximal firing along the corridor. **B**. Percentages of CA1 (*left*) or V1 (*right*) neurons with a significant spatial shift of their response across theta phases (‘*Theta-shifted neurons*’; p < 0.05), as a function of the position of maximal firing along the corridor. **C**. Examples of V1 neurons with a significant spatial shift of their response profiles across theta phases (p < 0.05). For each neuron, we plotted from top to bottom: *1)* the mean response profile; *2)* the raster plot of the theta phase of the spikes as a function of positions in the corridor; *3)* the mean firing rate as a function of theta phases and positions; *4)* the spatial cross-correlogram between the response profile at each theta phase and the mean response profile. **D**. Distribution of the amplitude of the spatial shift across theta phases for CA1 (*left*) or V1 (*right*) neurons whose responses were significantly shifted. The amplitude of the spatial shift was measured from the amplitude of the sinusoid that best fitted the maximum curve of the spatial cross-correlogram (*black curve* in **C**). Putative V1 interneurons were identified according to their spike waveform. Percentages are expressed relative to the total number of neurons within each category (CA1, n = 1422; V1 interneurons, n = 240; V1 pyramidal neurons, n = 869). **E**. Distribution of the linear-circular correlation coefficient measured from CA1 (*left*) or V1 (*right*) neurons. The linear-circular correlation coefficient was measured from the mean firing rate as a function of theta phases and positions (firing rate maps in **C**). *Black bars*: neurons which showed a significant linear-circular correlation (p < 0.05, CA1, 42.5%; V1, 11.45%); *gray bars*: all neurons.

